# A simple and efficient CRISPR technique for protein tagging

**DOI:** 10.1101/597971

**Authors:** Fanning Zeng, Valerie Beck, Sven Schuierer, Isabelle Garnier, Carole Manneville, Claudia Agarinis, Lapo Morelli, Lisa Quinn, Judith Knehr, Guglielmo Roma, Frederic Bassilana, Mark Nash

**Affiliations:** Novartis Institutes for Biomedical Research, Basel, Switzerland

## Abstract

Genetic knock-in using homology directed repair is an inefficient process, requiring selection of few modified cells and hindering its application to primary cells. Here we describe <u>**H**</u>omology <u>**i**</u>ndependent gene <u>**Tag**</u>ging (HiTag), a method to tag a protein of interest by CRISPR in up to 66% transfected cells with one single electroporation. The technique has proven effective in various cell types, can be used to knock in a fluorescent protein for live cell imaging, to modify the cellular location of a target protein and to monitor levels of a protein of interest by a luciferase assay in primary cells.

---

Genome editing has been revolutionized by the discovery of the CRISPR-Cas9 system. Cas9 nuclease creates a double-strand DNA break (DSB) at a target site specified by the guide RNA (gRNA) sequence^1,2^. In the presence of a donor DNA template, gene tagging can be achieved through homology-directed repair (HDR), the efficiency of which, however, is low and it might not work in all cell types. Compared with HDR, non-homologous end-joining (NHEJ) is more efficient and is active in both proliferating and post-mitotic cells. Indeed, DSBs are preferentially repaired via NHEJ pathway^3^. Although commonly used for gene knock-out by causing insertion or deletion (indel) mutations, NHEJ repair has been demonstrated to be intrinsically accurate and can be also used to create gene knock-ins^4-7^.

The application of CRISPR knock-in in cell biology is hampered by its low efficiency and laborious procedure. The aim of this study was to develop a CRISPR knock-in protocol that is: highly efficient; 2) easy to use; 3) applicable to cell lines and primary cells. We chose electroporation as the delivery method for gRNA/Cas9 nuclease/donor DNA complex, as it offers better genome editing efficiency ^8-10^. We opted to use dsDNA without any homology arms as the donor, and hypothesized that if dsDNA donors are available in abundance when and where DSB happens, they could be incorporated into the genome via NHEJ with high degrees of accuracy. With this strategy, we can efficiently tag and test the target proteins within a short timeframe. We name it <u>**H**</u>omology <u>**i**</u>ndependent gene <u>**Tag**</u>ging (HiTag).

To increase the chance of target integration, we limited the size of donor DNA to 100bp and thus focused initially on peptide tags. First, we aimed to label the heterogeneous nuclear ribonucleoprotein A2/B1 (HnRNP) in HeLa cells with a V5 tag. A gRNA was designed to target the 5’ UTR of *HNRNPA2B1*, and the dsDNA donor was synthesized as complementary oligonucleotides and annealed. To ensure that the *V5* tag is translated in frame, an ATG was added to its 5’ end, as well as 1bp to 3’ end (**Fig. 1a**). Three days after transfection, knock-in efficiency was assessed by anti-V5 immunostaining: 40±1.5% of the cells displayed strong V5 signals that exclusively localized in the nucleus (**Fig. 1b**). To further confirm the correct insertion of the V5 tag into the HnRNP, we performed western blotting to check the size of tagged proteins: two bands were visualized by the V5 antibody, with their sizes corresponding to the predicted chimeras (**Fig. 1c**). Although this result already suggested that the *V5* sequence was integrated into *HNRNPA2B1* as intended, we further confirmed its correct location at nucleotide precision by Amplicon sequencing (**Supplementary Fig. 1** and **2**).

**Figure 1.**
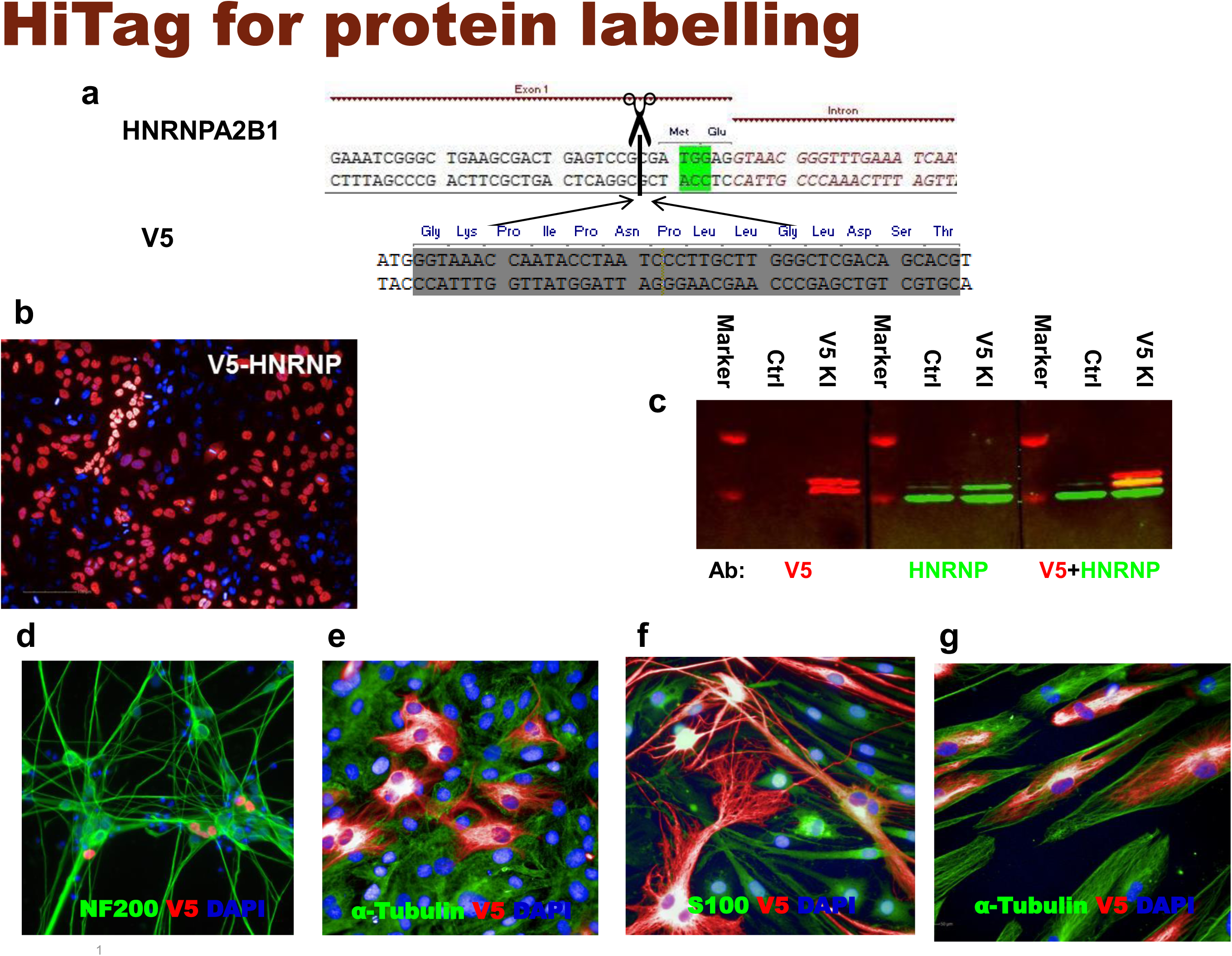
HiTag enables high efficient CRISPR knock-in for imaging. a. Sequences for *HNRNPA2B1* (5’ end) with the PAM site (green) and cleavage site (dotted line); V5 sequence with ATG and an extra base at the 3’
b. Immunostaining with V5 tagged HnRNP in HeLa cells (n=6)
c. Lysates from untransfected or V5-HnRNP knock-in HeLa cells were subjected to western blotting. Both V5 (left) and HNRNP (middle) antibody visualized two bands in western blots, corresponding to A2/B1 isoforms. V5 labelled fragments are slightly larger than those recognized by HnRNP antibody (right)
d. 5.9±1.6% iNgn2 cells display V5-HNRNP (red) signals. Cells were co-stained with neuronal marker NF200 (green) (n=4)
e. V5-Vimentin (red) in primary rat tenocytes, co-stained with α-Tubulin (green) (n=10)
f. V5-Vimentin (red) in primary rat SCs, co-stained with SC marker S100 (green) (n=8)
g. V5 knock-in Vimentin (red) in primary human skeleton muscle cells, co-stained with α-Tubulin (green) (n=4)

We investigated HiTag efficiency with several other targets in HeLa cells, such as β-actin, Vimentin and a cytosolic / nucleoli protein (RLP10A), all of which were successfully tagged (**Supplementary Fig. 3**). The method achieved similarly high efficiencies in other common cell lines (31.3∼50.7%) (**Supplementary Fig. 4**). We also achieved positive knock-in in human induced pluripotent stem (iPS) with the HiTag method (**Supplementary Fig. 5**).

CRISPR knock-in is generally not applicable to post-mitotic cells since their HDR mechanism is inactive. However, by using iPS derived iNgn2 neurons, we demonstrated that HiTag approach also worked in these cells (**Fig. 1d**). Primary cells can be problematic for CRISPR knock-in too, as they are not suitable for clonal selection. We extended the application of HiTag into primary cells: as shown in **Figs. 1e and f**, Vimentin can be labeled in 23.9±0.6% primary rat tenocytes, as well as 36.3±2.5% rat Schwann cells (SCs), while V5 tagged Sox10 was visualized in 9.9±0.2% cells (**Supplementary Fig. 6**). Surprisingly, HiTag protocol was even more efficient in the primary human skeleton muscle cells (SkMC): V5-Vimentin signals could be observed in 66.3±2.5% cells (**Fig. 1g**).

Gene tagging has applications in cell biology far beyond just labelling a protein for immunostaining. For example, an endogenous protein can be tracked in live cells by tagging with fluorescent proteins^5^. We introduced a dsDNA fragment encoding mScarlet^11^ into Vimentin or β-actin in HeLa cells via HiTag. Filament structures were observed in 0.6~1.5% live cells after 7 days (**Fig. 2a**), and mScarlet positive cells enriched by fluorescence-activated cell sorting. The subcellular location of mScarlet-Vimentin was then recorded in a time-lapse video (**Supplementary Video**).

**Figure 2.**
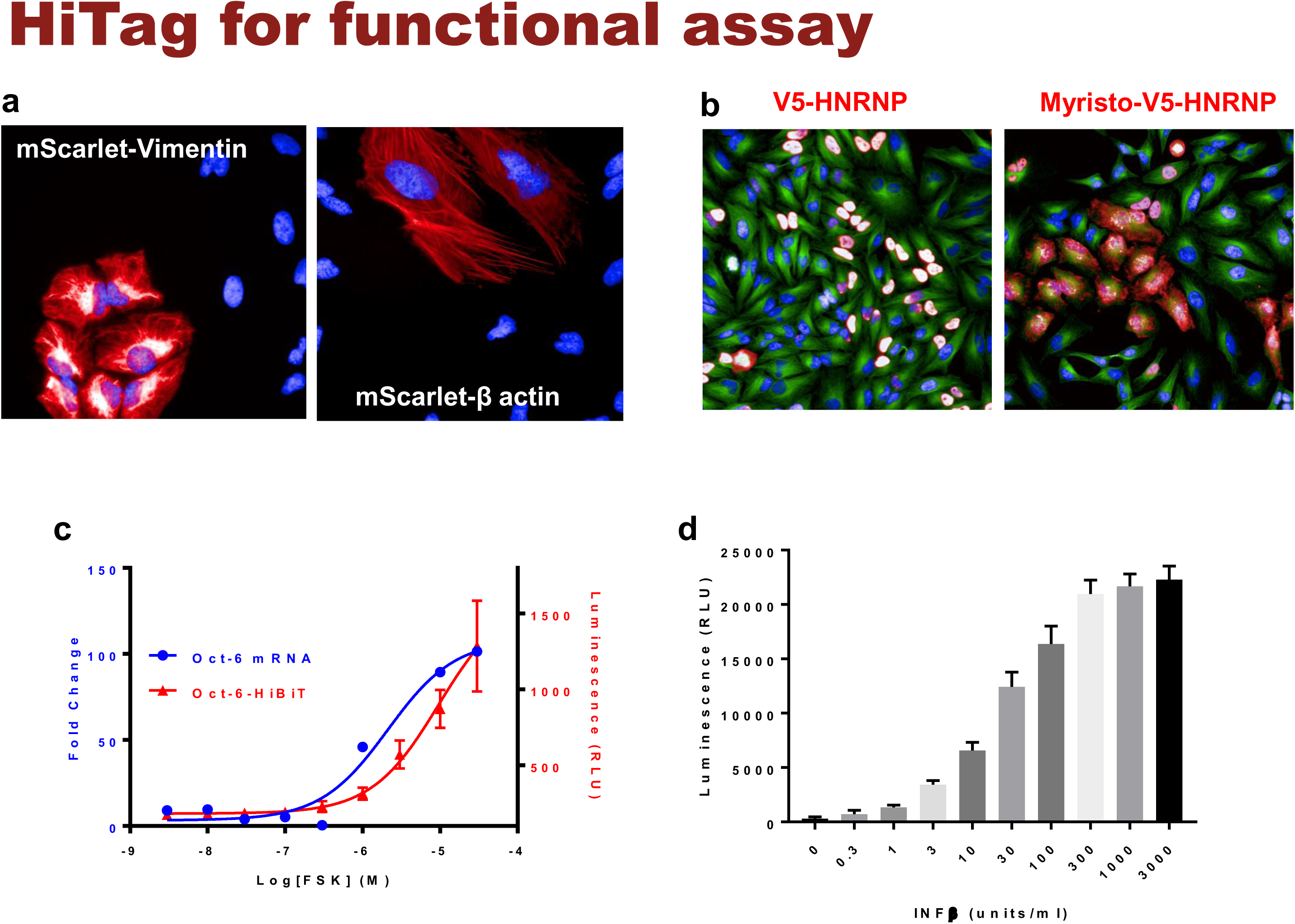
Multiple applications of HiTag in cell biology. a. mScarlet was inserted into the N-terminus of Vimentin or 5’ UTR of β-actin in HeLa cells. Live cell images were taken 7 days after transfection (n=10 each)
b. V5 HNRNP (left) and Myristoylated V5-HNRNP (right) in HELA cells. Addition of myristoylation sequence targets the endogenous protein to cell membrane (n=5)
c. Primary rat SCs were incubated with FSK for 24 hours, and Oct-6 mRNA expression were determined by qPCR (blue trace). To measure Oct-6 at the protein level, HiBiT sequence was inserted at its C-terminus (n=3). The cells were treated with FSK at various concentrations and subjected to NanoLuc assay after 48 hours (red trace) (n=4)
d. IFIT1 was tagged with HiBiT sequence at its C-terminus in primary human SkMC cells. The cells were treated with IFNβ at various concentrations and subjected to NanoLuc assay after 16 hours (n=10)

Another possible application is to change the cellular location of the target protein by attaching a signal peptide. As a proof of concept, a myristoylation sequence was added to V5-HnRNP (**Supplementary Table 1**) to disrupt its nucleus location: while 96.9±0.9% control cells display V5 signals in nuclei, 71.1±8.5% signals were cytosolic or on cell membranes when the myristoylation peptide was integrated (**Fig. 2b**).

Given the high efficiency of the HiTag protocol, we wondered whether it was possible to develop functional assays in primary cells via CRISPR knock-in. Oct-6 is a key transcription factor in the myelination pathway in Schwann cells. Its mRNA expression is up-regulated upon stimulation of cAMP pathway^12^ (**Fig. 2c**). To measure this up-regulation at the protein level, a HiBiT sequence (an 11 amino acid subunit of Nanoluc luciferase^13^) was attached to the C-terminus of Oct-6 protein in rat primary Schwann cells by HiTag. After the cells were incubated with Forskolin (FSK), which increases intracellular cAMP level, up-regulated Oct-6 protein expressions were evident in the Nanoluc assay (**Fig. 2c**).

To further validate this approach, HiBiT sequence was attached to the C-terminus of IFIT1 protein in human SkMC. After incubation with IFNβ, which induces IFIT1 expression^5^, a dose dependent response was observed in the luciferase assay (**Fig. 2d**). Therefore, we were able to develop a high throughput format assay to measure endogenous protein expression in primary cells via Crispr knock-in.

Here, we report HiTag as a simple and efficient protocol Crispr knock-in method that can be applied to various targets in a wide range of cell types. Although its exact mechanism remains elusive, HiTag knock-in seems to involve DNA-dependent protein kinase (DNA-PK), as the DNA-PK inhibitor^14^ abolished the knock-in effect. Unexpectedly, SCR7, a commonly used NHEJ pathway inhibitor^15^, did not block HiTag knock-in at all, suggesting DNA ligase IV might not be involved **(supplementary Fig. 7**).

After cutting of the genomic DNA by Cas9, knock-in of dsDNA donors need to compete with self-ligation of free ends. It is conceivable that the more dsDNA is used in transfection, the higher the knock-in efficiency would be. Indeed, that was shown in **Supplementary Fig. 8a**. Also, an inverse correlation between knock-in efficiency and the length of dsDNA donors is evident **(Supplementary Fig. 8b)**. In some cell types, we observed cell toxicity issues when 1uM or higher concentrations of donor DNAs were used, especially when they are more than 100bp in length (data not shown). Interestingly, although Cas9 nuclease (in RNP complex) is commonly used at 1~4 μM for electroporation (IDT website), we noticed that increased Cas9 concentration did not increase the knock-in efficiency (**Supplementary Fig. 8c**).

Compared with other CRISPR knock-in methods, one apparent advantage of HiTag method lies in its efficiency. While many published data claimed high knock-in efficiency, the numbers quoted were after selection or enrichment^5,16,17^. With HiTag protocol, however, we can achieve 30-60% efficiency (on the cellular level) without selection.

The second benefit of our protocol lies in its ease and simplicity: while other NHEJ based methods rely on vector expression of gRNA and Cas9 protein^5,6,16,17^, which requires intensive and expensive laboratory work, all reagents used in HiTag methods are readily available from commercial providers. In particular, we realize that synthetic dsDNAs are far more cost-effective donors compared to PCR amplicons or plasmid fragments. Combined with the fact that clonal selection is not necessary with HiTag, experiments on target proteins could be finished within two weeks, from design to a functional test.

Lastly, the importance of our method also lies in its diverse application. Primary cells are generally not suitable for Cripsr knock-in. However, given high efficiency of HiTag methods, we have established several luciferase assays to monitor the expression of endogenous proteins in primary cells.

Like other CRISPR knock-in methods, the successful application of HiTag depends on the targeting and cutting efficiency of gRNA/Cas9 complex; and the detection of the tagged proteins heavily relies on their endogenous expression level. HiTag approach has also its own intrinsic limitation: the orientations of the integrated donor DNAs are non-controllable with Cas9 nuclease. Furthermore, the delivery of RNP into cells via electroporation is essential for HiTag, thus its throughput is rather low in the current format. However, electroporation devices are available in 96 well format as the alternative. Nevertheless, HiTag offers a highly efficient CRISPR knock-in strategy that can be applied to many target genes and cell types. We expect the simplicity of this approach will encourage more studies to be performed on the endogenous proteins in their native environments.

## Methods

Methods and any associated accession codes and references are available in the online version of the paper.

---

**Supplementary figure 1:**
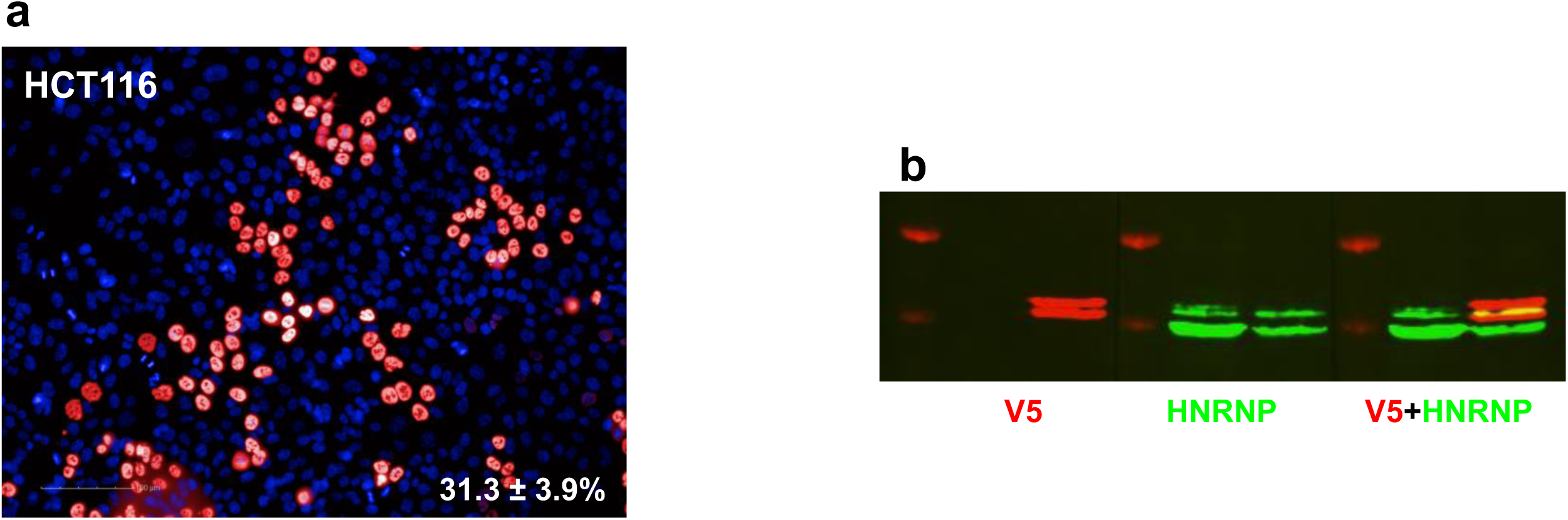
HiTag enables high efficiency CRISPR knock-in of HCT116 cells. The diploid HCT116 cell line has been favored for the parallel analysis of KI efficiency by Next Generation Sequencing (NGS) because of its more stable chromosomal composition. Therefore, V5 tag knock-in in HnRNP was performed in HCT116 cells.

a. 3 days after transfection, immunostaining with V5 antibody shows that 31.3 ± 3.9% HCT116 cells display positive signals in nucleus (n=5).
b. Lysates from untransfected or V5-HnRNP knock-in HCT116 cells were subjected to western blotting. A2/B1 isoforms can be detected with either V5 (left) or HNRNP (middle) antibody, and with V5+HNRNP co-staining (right)

**Supplementary figure 2:**
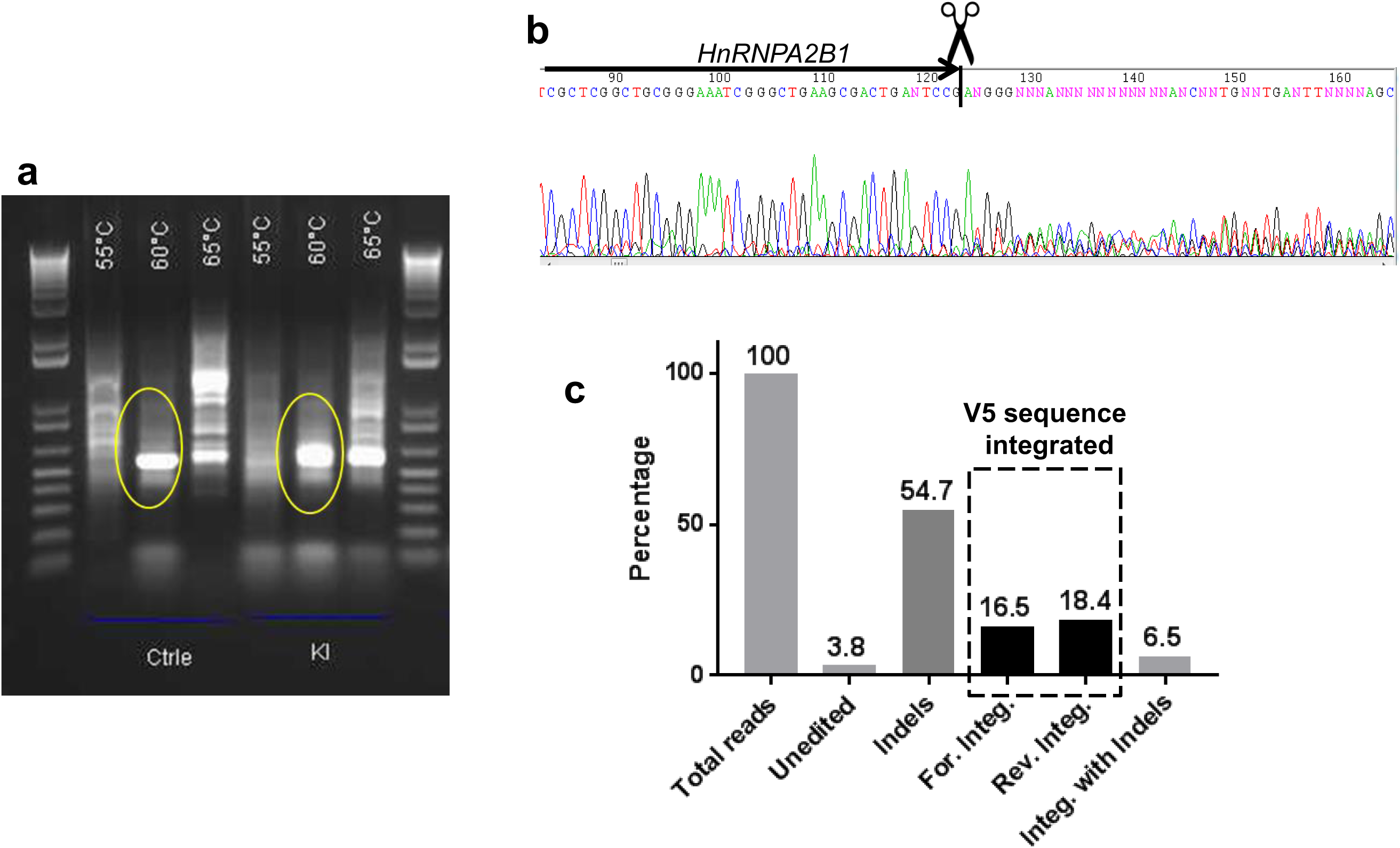
Amplicon sequencing result in HCT116 cells. a. The region of theoretical insertion was amplified from genomic DNA with various PCR parameters. The circled PCR products were purified and submitted for Sanger and NGS sequencing.
b. Sanger sequencing result clearly suggests editing events after Cas9 cleavage site.
c. The results from Amplicon-sequencing showing percentage of fragments falling into different alignment categories: Unedited fragments align to the wildtype genome and contain no indels whereas Indels fragments also align to the wildtype genome but contain at least one indel. Forward/Reverse Integrated fragments align to the genome sequence containing the 46bp insert between bases 26200579 and 26200580 on chromosome 7 without an indel. The classification of the direction of the alignment as forward or reverse is performed regarding the orientation of the gene HNRPA2B1 – which is located on the reverse strand of chromosome 7. Integrated with Indels fragments also align to the genome sequence containing the 46bp insert but contain an indel. As expected, the ratio of forward and reverse integration of V5 sequence is about 50:50, and only forward integration (16.5%) can result in a functional expression of V5 tag. To calculate functional V5 tagging at cellular level, we use the following equation:

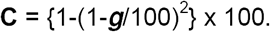 ***g*** stands for percentage of functional **g**enome integration, which is 16.5 in this case; **c** represents the percentage of functional V5 expression at **c**ellular level, which is 30.3. The number matches well with the immunostaining result (Supplementary Fig. 1), which suggested 31.3 ± 3.9% cells express V5-HnRNP.

**Supplementary figure 3:**
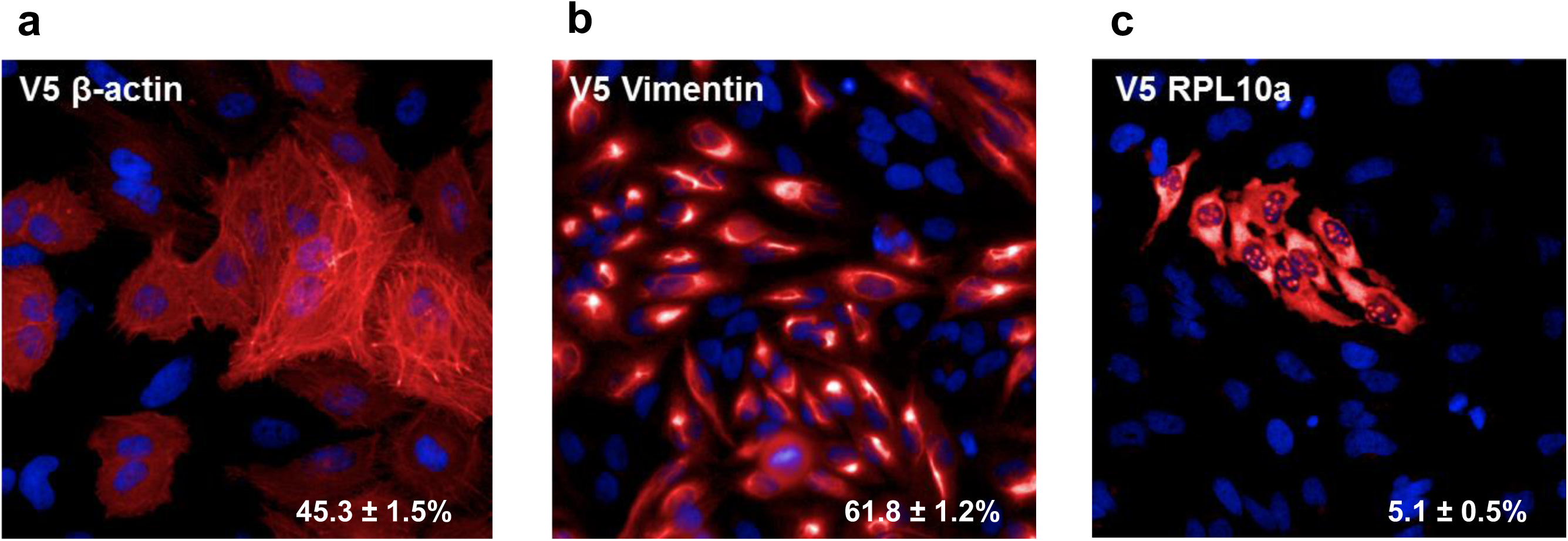
V5 tag knock-in in β-actin, Vimentin and RLP10a in HeLa cells. a. 3 days after transfection, 45.3 ± 1.5% cells display actin filament structures after staining with V5 antibody (n=5)
b. After V5 knock-in in Vimentin, intermediate filament structures were observed in 61.8 ± 1.2% cells with V5 antibody staining (n=5)
c. V5 sequence was knocked into *RPL10A*, which encodes Ribosomal Protein L10a. Cells were stained with V5 antibody: 5.1 ± 0.5% cells display positive signals in cytosol and nucleoli (n=5)

**Supplementary figure 4:**
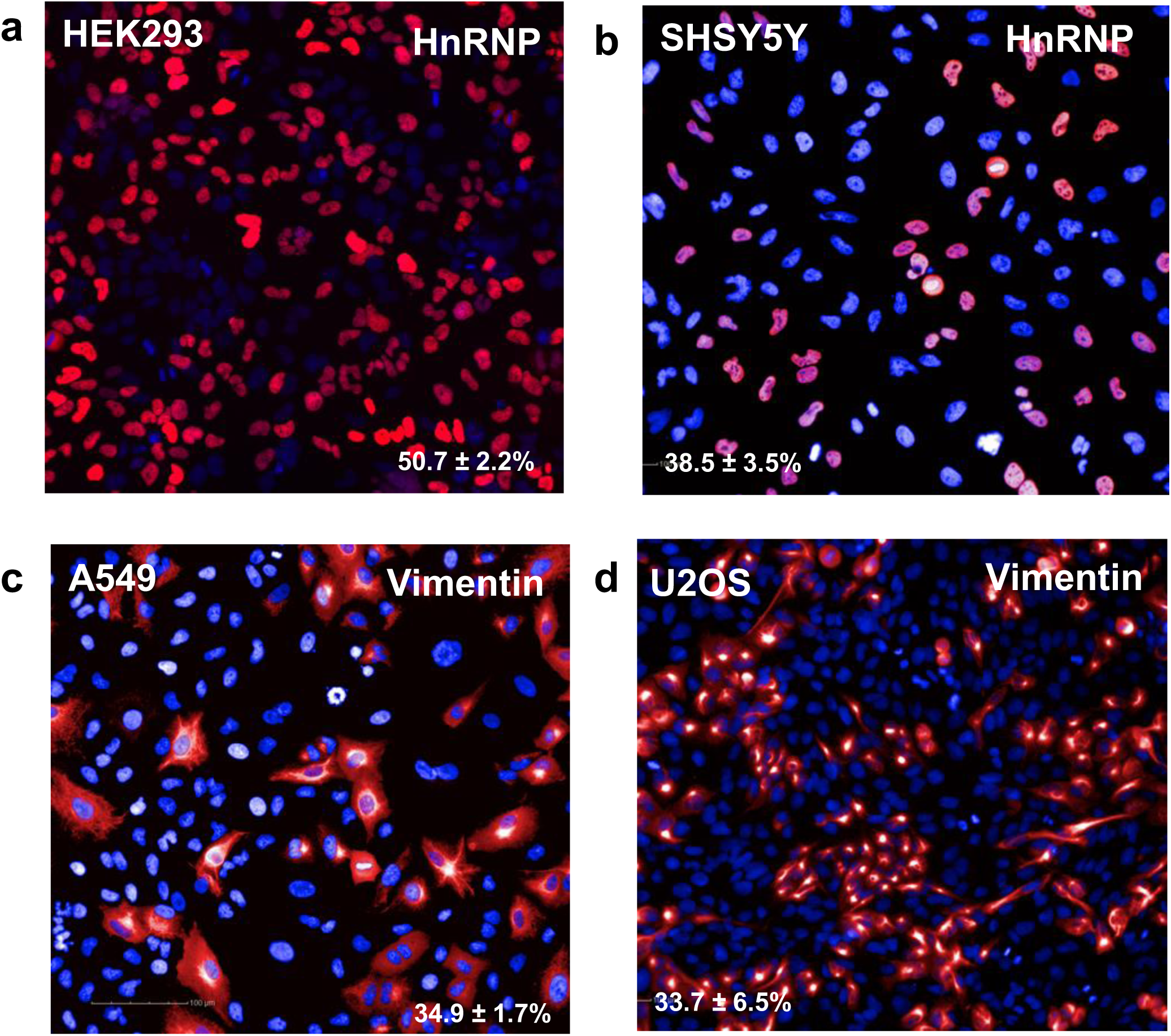
HiTag efficiency in other common cell lines. a. For *HNRNPA2B1* in HEK293 cells, V5 signals were detected in 50.7 ± 2.2% cells after CRISPR knock-in (n=6)
b. After V5 knock-in via HiTag approach, 38.5 ± 3.5% SH-SY5Ycells display V5 signals in nucleus (HnRNPA2B1) (n=4)
c. V5 knock-in with HiTag can achieve 34.9 ± 1.7% efficiency in Vimentin in A549 cells (n=3)
d. With U2OS cells, knock-in efficiency was 33.7 ± 6.5% in Vimentin (n=6)

**Supplementary figure 5:**
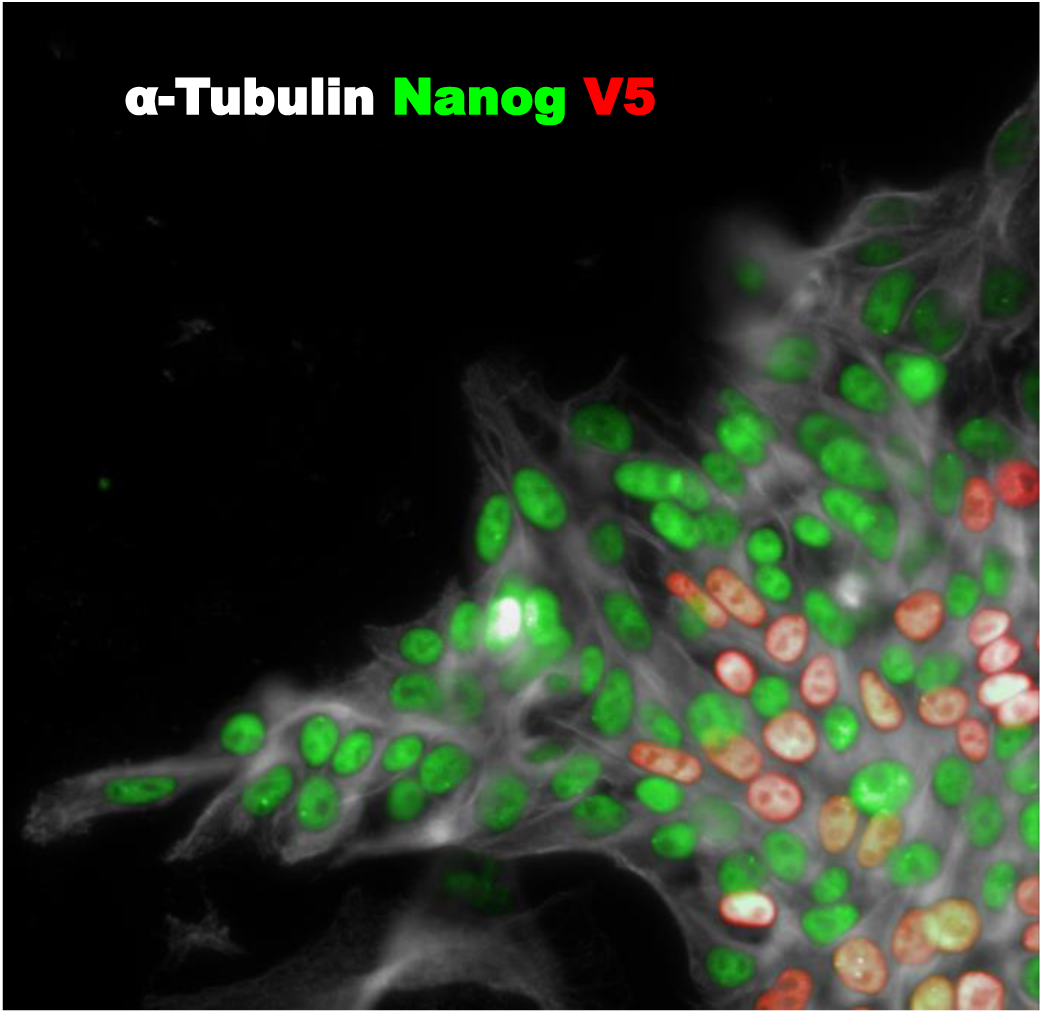
HiTag efficiency iPS cells. V5 sequence was targeted into HnRNPA2B in human iPS cells. 3 days after electroporation, V5 knock-in (red) was achieved in 10.4±1.8% iPS cells. Cells were co-stained with pluripotency marker Nanog (green) and α-Tubulin (grey) (n=5)

**Supplementary figure 6:**
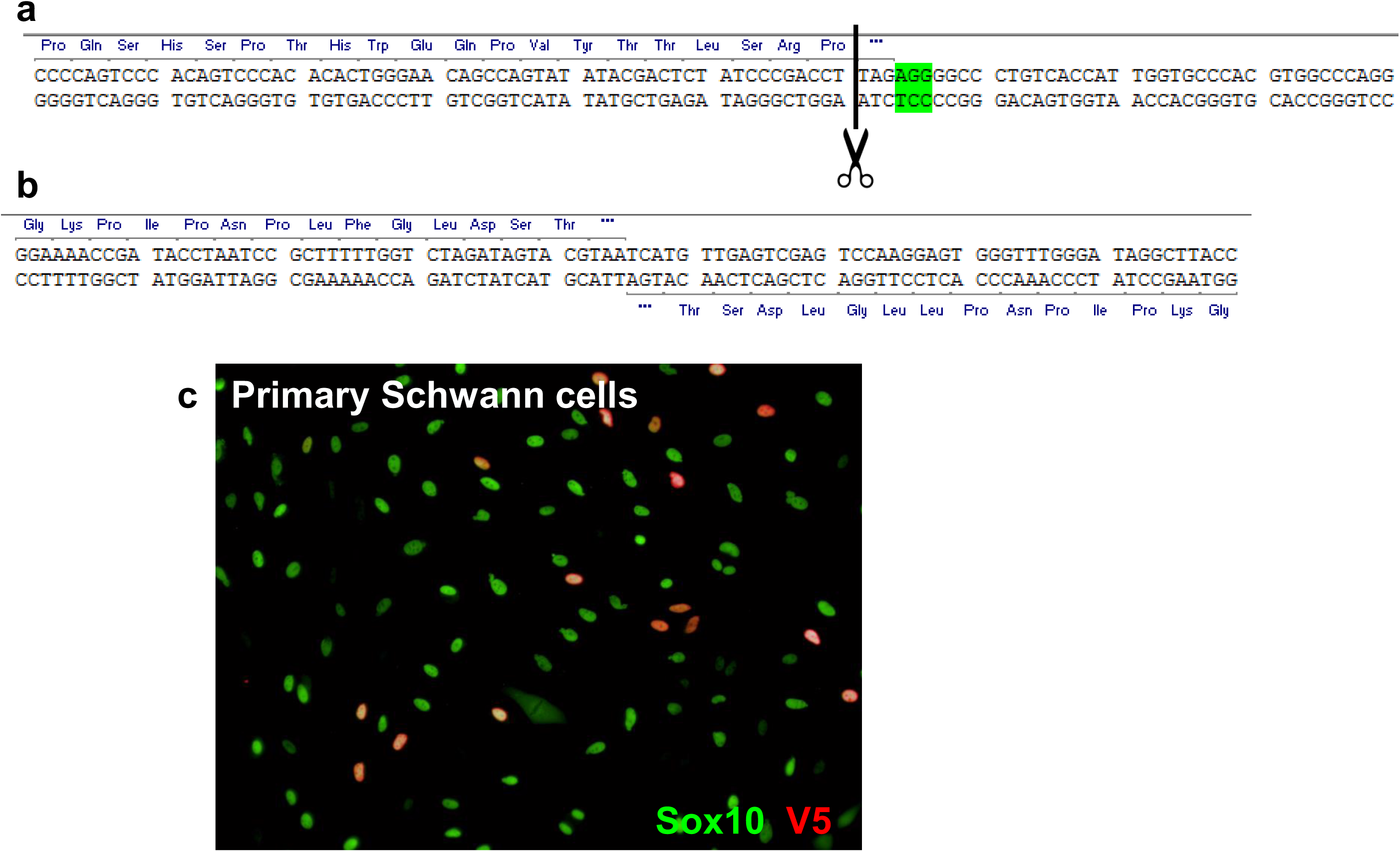
HiTag efficiency in primary Schwann cells. V5 sequence was targeted into *SOX10* gene (which encodes a transcription factor) in rat primary Schwann cells.

a. Genomic sequence for rat *SOX10* (at 3’ terminus of its CDS). PAM site is highlighted in green and Cas9 cutting site is indicated with a dotted line
b. To increase the chance of functional V5 tag integration, a dsDNA donor containing the V5 sequence and a stop codon in both directions was used
c. CRISPR knock-in was performed via HiTag approach on rat primary Schwann cells. Efficiency was assessed by immunostaining with V5 (red) and Sox10 (green) antibodies (n=4)

**Supplementary figure 7:**
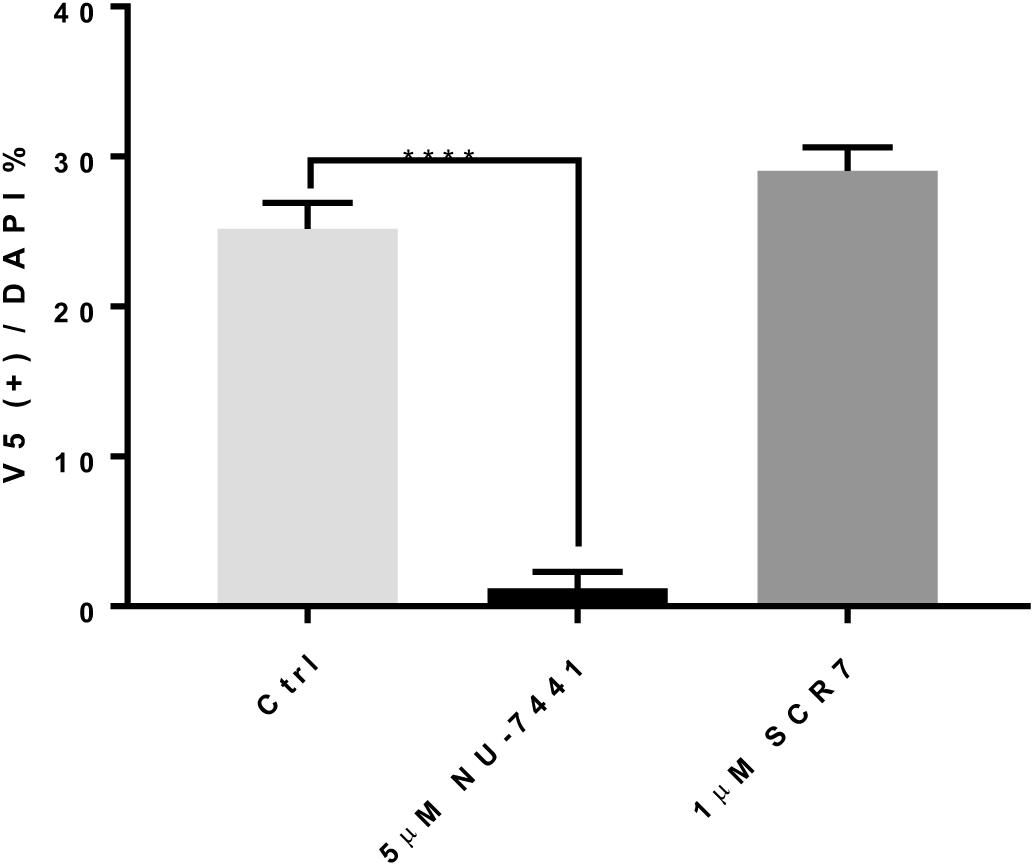
Effect of NU-7441 and SCR7 on HiTag knock-in efficiency. HeLa cells were treated with 5μM NU-7441 and 1μM SCR7 for 2 and 24 hours respectively prior to the transfection. The V5 sequence was knocked into *HNRNPA2B1*, and its efficiency was assessed after 72 hours by immunostaining (n=8)

**Supplementary figure 8:**
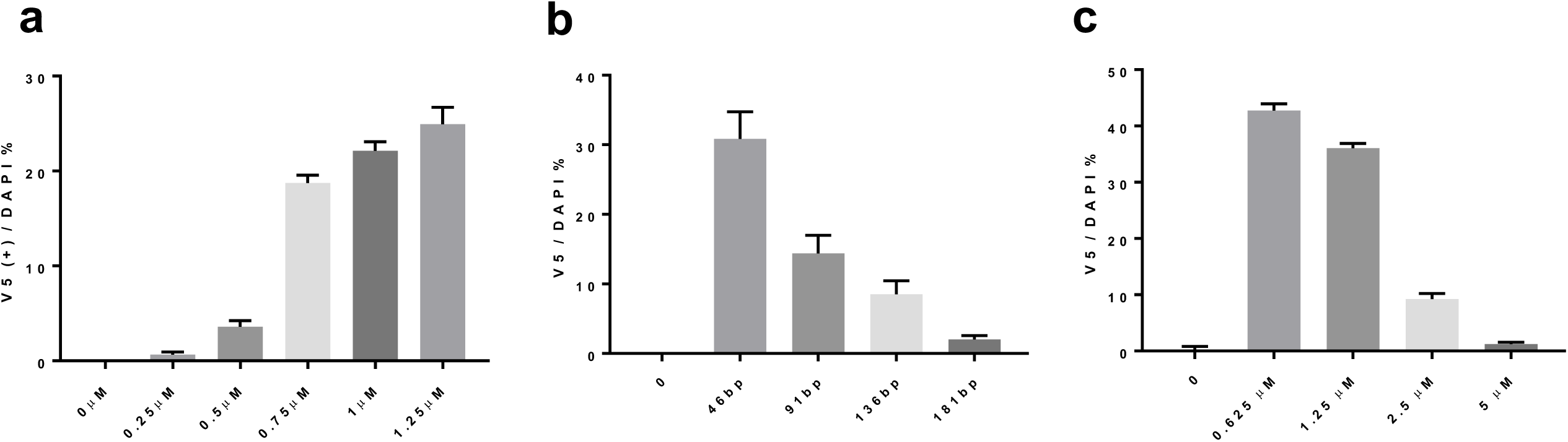
Effect of various parameters on HiTag knock-in efficiency. By targeting *HNRNPA2B1* in HeLa cells, we assessed the effects of several parameters on the knock-in efficiency of V5 sequence.

a. 46bp donor DNAs containing V5 sequence was knocked into *HNRNPA2B1*, with dsDNAs used at various concentrations during electroporation. Cells were stained with V5 antibody after 72 hours (n=8)
b. Donor DNAs containing one copy V5 sequence and linkers at various length (Supplementary Table 1) were knocked into *HNRNPA2B1*. Cells were stained with V5 antibody after 72 hours to assess knock-in efficiency (n=8)
c. 46bp donor DNAs containing V5 sequence was knocked into *HNRNPA2B1*, with various concentrations of Cas9 nuclease being used during electroporation. Cells were stained with V5 antibody after 72 hours (n=8)

## Supplementary video 1

An mScarlet encoding sequence lacking start and stop codons was knocked in *VIM* gene in HeLa cells by HiTag protocol. 9 days after transfection, the cells were subjected to FACS sorting, and mScarlet positive cells were collected and cultured for further 4 days. For live cell imaging, cells were seeded in a 96 well plate and time lapse images were taken over a 16 hour period on Yokogawa CV7000 imaging system. It is evident that cellular distribution of mScarlet-Vimentin can change from filament like structures to punctate appearance.

## Methods

### CRISPR knock-in

#### gRNAs

we used Alt-R CRISPR-Cas9 System from Integrated DNA Technologies (IDT). The gRNAs were designed using CRISPOR program^18^ (http://crispor.tefor.net/) and listed in Supplementary Table 2. crRNA and tracrRNA were dissolved with Duplex buffer (IDT) to 169μM each. They were mixed at the equal volume, and the final RNA concentration adjusted to 2 μg/μl by adding Duplex buffer. gRNA complex was formed by incubation at 95°C for 5 minutes, followed slowly cooling down to room temperature (RT), then stored at −80°C.

#### Cas9 nuclease

Cas9 proteins were mostly produced in house. Briefly, the NLS-SpyCAS9-NLS-His6 was expressed in E. coli strain Rosetta 2 (DE3) in pET28a+, using 2xYT media plus 50 μg/ml kanamycin overnight at 18°C. The Cas9 was then purified using Ni-NTA agarose (Qiagen) capture/elution, followed by size exclusion chromatography on a S200 26/600 Superdex column (GE Lifesciences) using modified buffers^19^ from Jinek et al. Then Cas9 stocks (5.9 mg/ml) were aliquoted and stored at −80°C. Commercially available Cas9 nuclease (IDT) can be also used, with no difference observed in our hands (data not shown). When HiTag protocol was tested in HCT116 cells stably expressing Cas9, the knock-in efficiency is equivalent to that by Cas9 proteins (data not shown).

#### dsDNA donors

Oligonucleotides were synthesized by Microsynth with PAGE purification. The forward and reverse strands were dissolved in Duplex buffer to 20 μM each. To anneal the oligos, they were mixed at equal volume and incubated at 95°C for 5min, then were allowed to cool down slowly to RT. The final concentration of dsDNA stock is 10μM. Unless stated otherwise, final concentrations of dsDNA in the transfection mixtures are 750 nM.

#### Transfection mixture

For each transfection, we prepared 20 μl of cell / ribonucleoprotein / dsDNA mixture. Cells were trypsinized and washed once with phosphate-buffered saline (PBS), then resuspended at the concentration of 5.4.×10^6^ cells per 100 μl buffer R (IDT). Ribonucleoprotein (RNP) was formed by incubating 4 μg Cas9 (0.68 μl), 2 μg gRNA (1μl) at RT for 10min in a final volume of 5 μl Duplex buffer. Then 400,000 cells (in 7.5 μl buffer R), 15 pmol dsDNA (1.5 μl) and 6 μl buffer R were added to make up the transfection mixtures (Supplementary Table 3).

For knock-in of mScarlet, dsDNA donors were ordered as gBLOCK gene fragments (IDT). 500ng DNA was dissolved in 5 μl buffer R (IDT). The constitution of transfection mixtures are listed in Supplementary Table 3.

#### Electroporation

The RNP and dsDNA template were delivered into cells by electroporation with Neon Transfection System (ThermoFisher, MPK5000). We used the 10 μl kit (MPK1096), and transfection protocols for individual cell type were listed in Supplementary Table 4. After transfection, cells were cultured in antibiotic free medium for at least 24 hours before replacing with normal growth medium.

### Cell culture

All cells were maintained under a humidified atmosphere of 5% CO_2_ at 37°C. A549, HCT116, HEK293, HeLa, SH-SY5Y and U-2 OS cell lines are purchased from ATCC and cultured according to supplier’s recommendation. The primary human skeletal muscle cells (Lonza, CC-2561) were maintained in Skeletal Muscle Growth Medium (Lonza, CC-3246), supplemented with 20% fetal calf serum (FCS).

#### Schwann cells culture

Primary Schwann cells were isolated as described previously by Kaewkhaw *et. al*.^20^ Briefly, Sciatic nerves were isolated from adult rats and epineurium stripped. The tissues were teased with sharp tweezers, and digested with 0.05% collagenase digestion at 37°C for 2 hours. Homogenous cell suspension was obtained by dissociating the tissue with glass Pasteur pipette and filtered through 40 μm cell strainer. The cells were pelleted down and plated in poly-D-lysine coated flasks. The cells were maintained in a custom made D-valine DMEM medium (based on 10313039, L-Valine free, ThermoFisher), supplemented with 10% FCS, B27, 10mM Hepes, 2mM glutamate and 5μM Forskolin. Once the cultures reach confluency, Schwann cells were dispatched by Dispase (Corning) digestion, and can be sub-cultured for up to 10 passages. The purities of Schwann cell were at least 95% at the time of transfection, judged by immunostaining with Schwann cell markers.

#### Generation and maintenance of iPS cells

Neonatal human dermal fibroblasts (ThermoFisher) were used for reprogramming using Sendai virus with the help of the CytoTune-iPS reprogramming kit according to the standard protocol. Colonies with hallmark of pluripotent morphology were readily visible between days 17 and 20 after transduction. These were picked and sub-cloned multiple times on plates coated with Matrigel (BD Biosciences) in mTeSR1 medium (STEMCELL Technologies) until Sendai virus RNA could no longer be detected and the morphology looked stable. Pluripotency was controlled by FACS analysis. Potential to differentiation into the three germlayers was approved by using the TaqMan hPSC Scorecard Panel (ThermoFisher) according to the supplier guideline. Karyotype analysis was performed by full-genome SNP analyses done by Life & Brain and showed no larger chromosomal aberrations. iPSC were maintained on matrigel-coated plates and grown in mTesR1.

#### Generation and differentiation of iNgn2 iPSCs

Human Ngn2 cDNA was synthesized using sequence information from the Ensembl database (Ensembl Gene ID ENSG00000178403) and cloned under the control of TRE tight (Tetracycline Response Element) promoter in a PiggyBac/Tet-ON all–in-one vector^21^. This vector contains a CAG rtTA16 cassette allowing constitutive expression of Tet-ON system and an Hsv-tkNeo cassette for generation of stable IPS clones. After trypsinization into single cells with Tryple express reagent (ThermoFisher) approximately 1 × 10^6^ iPS cells were nucleofected by Amaxa nuclefector device using Human Stem Cell Nucleofector Kit 1 (Lonza) and Program B-016 with 4 μg of Ngn2 plasmid and 1 μg of the dual helper plasmid. Subsequently cells were replated on matrigel plates with mTeSR medium containing 10 μM of Rock inhibitor. Antibiotic selection (G418 0.1 mg/ml) was applied 48 hours later. Stable clones appear within 1 week. To differentiate iNgn2 neurons, 1 × 10^6^ of iPS cells were plated on a 6 cm matrigel plate in proliferation medium (DMEM/F12 with Glutamax supplemented with 2% B27 and 1% N2, 10 ng/ml hEGF, 10 ng/ml hFGF, 1% Pen/Strep (all from ThermoFisher) containing Rock inhibitor (10μM) for 1d and doxycycline (1ug/ml) for 3d. Three days later cells induced neurons were frozen down. Cells were thawed and recovered in differentiation medium for 24 hours before transfection (Neurobasal supplemented with 2% B27, 1% N2, Pen/Strep, 1 mM Sodium Pyruvate, plus 10 ng/ml BDNF, GDNF and hNT3). To ensure the cells were post-mitotic at the time of electroporation, the cultures were treated with 10μM cytosine arabinoside 24 hours before and after.

### Amplicon sequencing

Total genomic DNA was isolated from transfected cells using a DNA extraction kit (Qiagen, DNeasy Blood and Tissue kit). The region of theoretical insertion was amplified by PCR using primers hnRNPA2B1 fw (5’-tcccgtgcggaggtgctcctcgcag) and hnRNPA2B1 re (5’-agctccgcagcctcgctcacgagg). A 500bp fragment could be obtained after 40 amplification cycles with denaturation 95°C for 30s - annealing 60°C for 30s – extension 68°C for 45s using a proofreading Accuprime Pfx Mix (ThermoFisher). After gel purification, the PCR fragments were used to prepare sequencing libraries with the Ovation Low complexity Sequencing System (NuGEN, cat.9092-256). Libraries were sequenced in paired-end mode, at a read length of 2 × 300 bp, using the MiSeq platform (Illumina). We obtained 10253514 and 11140907 read pairs for the control and the CRISPR samples, respectively. Read quality was assessed by running FastQC (version 0.10) on the FASTQ files. Since the sequencing quality decreased continuously after base 200 we trimmed the reads to a length of 240 base pairs before alignment. The FASTQ files were aligned against an extended human reference genome (based on GRCh38) using BWA version 0.7.15 with default parameter settings^22^. The extension of the genome consisted of the addition of two artificial chromosomes which contained the insert in forward and reverse orientation concatenated with 220 bp of chromosome 7 before base 26200579 and 220 bp after base 26200580. The alignments were then processed by a custom script to compute the different alignment classes of fragments.

### Western blot analysis

72 hours after transfection, cells were washed with PBS and lysed in RIPA buffer (ThermoFisher) containing a protease inhibitor cocktail (Roche). Protein concentrations in the samples were determined using a colorimetric assay based on the Lowry method (DC Protein Assay, Biorad) following manufacturer’s instructions. Equal amounts (ranging from 5 to 10ug) of total protein extracts were loaded after a 10mn denaturation step at 70°C on Nu-PAGE 4-12% Bis-Tris Gels (ThermoFisher) and separated by electrophoresis using MOPS buffer under reducing conditions. Proteins were transferred on a 0.2 μm PVDF membrane by electroblotting (TransBlot Turbo system, BioRad). After one hour room temperature blocking of the membrane in Odyssey PBS-Blocking buffer (LI-COR), the commercial primary antibodies anti-V5 (ThermoFisher), anti-Vimentin (Abcam), anti hnRNPA2B1 (Abcam) were diluted 1:2000 in fresh blocking buffer, added on the membrane and incubated overnight at 4°C. Membranes were washed twice in TBS buffer containing 0.1% Tween 20 (TBST), and incubated with fluorescence labelled secondary antibodies (Goat anti Mouse IR Dye 680 RD, or Goat anti-Rabbit IR Dye 800CW, LI-COR) diluted 1:10000 for 1h at room temperature. After 3 wash steps in TBST, the labelled proteins were visualized using an Odyssey Infrared Imaging system (LI-COR), and quantification of the obtained bands was done using the embedded Odyssey application software.

### Immunostaining, imaging and FACS sorting

Immunostaining was performed according to standard protocols. Briefly, 72 hours after transfection, cells grown on 96 well plates were fixed and permeabilized for 20 mins at 4°C with Fixation/Permeabilization solution (BD Biosciences). Cells were washed 3 times in PBS, then blocked in PBS containing 5% donkey serum, 1 % bovine serum albumin (BSA) and 10% Perm/Wash buffer (BD Biosciences) at RT for 1 hour. For V5 tag staining, cells were incubated at RT for 4 hours with Alexa647 conjugated anti-V5 antibody, followed by rinsing 3 times in PBS before imaging. For other antigens, cells were incubated overnight at 4°C with primary antibodies as indicated (Supplementary Table 5), washed three times with PBS, then incubated with 1:1000 Alexa Fluor-conjugated secondary antibodies (ThermoFisher) for 1 hour at RT. The Nuclei were visualized by counterstaining with NucBlue (ThermoFisher). Finally, the cells were rinsed a further 3 times in PBS before reading the plate on Operetta Imaging System (PerkinElmer). The image analyses were done with the built-in Harmony software (PerkinElmer).

For live cell imaging of mScarlet-Actin / Vimentin, HeLa cells were cultured on a 96 well plate for 7 days after transfection. The nuclei were stained NucBlue Live ReadyProbe (ThermoFisher) and the culture media replaced with FluoroBrite DMEM (ThermoFisher) supplemented with 3% FCS. The images were taken on Operetta and analysed with Harmony software.

To enrich mScarlet positive cells, HeLa cells were subjected to FACS sorting by standard protocol. Briefly, cells were trypsinized and resuspended in PBS without Ca^2+^ /Mg^2+^, with 2% FCS and 2 mM EDTA to obtain the single cell solutions (1×10^6^ cells/ml). Cells were sorted on BD FACSAria Fusion, with 561nm laser for excitation and a 610/20nm filter for emission. Positive cells were collected in a 6 well plate within the normal HeLa cell growth medium. Once the cells were confluent, they were split into a 96 well plate in FluoroBrite DMEM plus 3% FCS. Images were collected on Cell Voyager 7000 (Yokogawa) over 16 hours, with one image every 10 min. The movie was generated with ImageJ.

### Reverse Transcription Quantitative PCR

Total RNA was extracted from rat primary Schwann cells at the indicated time points with RNeasy (Qiagen) according to the manufacturer’s instructions. RNA concentration was determined by using a Nanodrop and reverse transcription was done with 300 ng of total RNA by Reverse Transcription kit (Thermo Fisher Scientific). Quantitative PCR (qPCR) reactions were carried out on a 7900HT AB instrument with TaqMan universal buffer (Thermo Fisher Scientific) and primers master mix designed in house and produce by Microsynth (Supplementary table 6). mRNA levels were normalized to levels of PpiB and ActB mRNA (Taqman probe from Thermo Fisher Scientific) in each sample.

### HiBiT Nano-Glo assays

A palindrome-like HiBiT sequence (supplementary table 1) was knocked into Oct-6 in rat SCs, and cells can be expanded for further 5~6 passages before use. The cells were split into a 384 well plate with 10000 cells per well. Cells were cultured for 3 days in FSK free medium, before FSK were added to the wells at various concentrations. The expression level of Oct-6-HiBiT was determined with the NanoGlo HiBiT lytic assay kit (Promega, N3040). Briefly, Nano-Glo substrate and LgBiT protein were mixed with the lytic buffer (1:50 and 1:100 respectively). The volume of culture media was adjusted to 30 μl each well, then equal volume of reagents was added. After 10 minutes, luminescence signals were measured on EnVision Plate Reader (PerkinElmer).

A palindrome-like HiBiT sequence (supplementary table 1) was knocked into IFIT1 in SkMCs, and cells expanded for 10 days in the growth medium. The cells were split into a 384 well plate with 5000 cells per well. After cells attached, Interferon-β (Merck, IF014) were added to the wells at various concentrations. The expression level of IFIT1-HiBiT was determined with NanoGlo HiBiT lytic assay kit.

### Statistical analysis

All statistical analyses were conducted using the GraphPad Prism. Data were expressed as the mean ± standard deviation.

## References

1. Cong, L. et al. Science 339, 819–823 (2013).

2. Hsu, P.D., Lander, E.S. & Zhang, F. Cell 157, 1262–1278 (2014).

3. Mao, Z., Bozzella, M., Seluanov, A. & Gorbunova, V. DNA Repair (Amst) 7, 1765–1771 (2008).

4. Suzuki, K. et al. Nature 540, 144–149 (2016).

5. Lackner, D.H. et al. Nat Commun 6, 10237 (2015).

6. He, X. et al. Nucleic Acids Res 44, e85 (2016).

7. Fei, J.F. et al. Proc Natl Acad Sci U S A (2017).

8. Leonetti, M.D., Sekine, S., Kamiyama, D., Weissman, J.S. & Huang, B. Proc Natl Acad Sci U S A 113, E3501–3508 (2016).

9. Liang, X., Potter, J., Kumar, S., Ravinder, N. & Chesnut, J.D. J Biotechnol 241, 136–146 (2017).

10. Burger, A. et al. Development 143, 2025–2037 (2016).

11. Bindels, D.S. et al. Nat Methods 14, 53–56 (2017).

12. Kawasaki, T., Oka, N., Tachibana, H., Akiguchi, I. & Shibasaki, H. Acta Neuropathol 105, 203–208 (2003).

13. Dixon, A.S. et al. ACS Chem Biol 11, 400–408 (2016).

14. Robert, F., Barbeau, M., Ethier, S., Dostie, J. & Pelletier, J. Genome Med 7, 93 (2015).

15. Maruyama, T. et al. Nat Biotechnol 33, 538–542 (2015).

16. Schmid-Burgk, J.L., Honing, K., Ebert, T.S. & Hornung, V. Nat Commun 7, 12338 (2016).

17. Sawatsubashi, S., Joko, Y., Fukumoto, S., Matsumoto, T. & Sugano, S.S. Sci Rep 8, 593 (2018).

## References

18. Haeussler, M. et al. Genome Biol 17, 148 (2016).

19. Jinek, M. et al. Science 337, 816–821 (2012).

20. Kaewkhaw, R., Scutt, A.M. & Haycock, J.W. Nat Protoc 7, 1996–2004 (2012).

21. Lacoste, A., Berenshteyn, F. & Brivanlou, A.H. Cell Stem Cell 5, 332–342 (2009).

22. Li, H. & Durbin, R. Bioinformatics 25, 1754–1760 (2009).

